# The RAE1-STOP1 Module Reduces Sensitivity to Exogenous ABA Treatment by Regulating ABI5 in Arabidopsis

**DOI:** 10.1101/2024.06.09.598107

**Authors:** Yuqing Zhang, Min Huang, Yinyin Liu, Mengmeng Yang, Yuqi Hou, Chao-Feng Huang, Ningning Wang, Lei Li

## Abstract

The SENSITIVE TO PROTON RHIZOTOXICITY 1 (STOP1) transcription factor plays a pivotal role in maintaining cellular ion balance and governing aluminum tolerance in plants. Abscisic acid (ABA) participates in aluminum tolerance by inducing the expression of several genes that are STOP1 targets. However, the mechanisms underlying ABA signaling and STOP1-mediated gene expression remain poorly understood. The F-box protein RAE1, an SCF-type E3 ligase component, recognizes STOP1 and controls its ubiquitination and degradation. This study revealed that exogenous ABA supplementation reduced STOP1 levels by promoting the expression of *RAE1*. Notably, both *RAE1* loss-of-function mutants and *STOP1* overexpressing lines showed enhanced sensitivity to exogenous ABA treatment, which correlated with early stage post-transcriptional upregulation of ABSCISIC ACID INSENSITIVE5 (ABI5). Our observations strongly suggest that RAE1 operates as an ABA-responsive factor, exerting control over STOP1 homeostasis, and thus establishing a negative feedback loop that controls ABA responses in Arabidopsis. Thus, our study revealed a novel function of the RAE1-STOP1 module in ABA signaling, highlighting its role in reducing ABA sensitivity by preventing ABI5 increase.

**Single Sentence Summary:** F-box protein RAE1 functions as an exogenous ABA responsive mediator to reduce STOP1-mediated ABA sensitivity.

## Introduction

The SENSITIVE TO PROTON RHIZOTOXICITY (STOP1) transcription factor is a key regulator of maintain cellular ion homeostasis (Balzergue *et al*., 2017; Sadhukhan *et al*., 2019; Sawaki *et al*., 2009). STOP1 can regulate cellular organic acid and ion status by activating the expression of *Aluminum-activated Malate Transporter* (*ALMT1*), *CALCINEURIN B-LIKE INTERACTING PROTEIN KINASE23* (*CIPK23*) and *GLUTAMATE DEHYDROGENASE* (*GDHs*) which participate in malate secretion, ion transport and pH regulation-associated metabolism (Sadhukhan *et al*., 2019; Sadhukhan *et al*., 2021). Specifically, STOP1 directly binds to the promoter of *nitrate transporter* (*NRT1.1*), promoting its transcriptional activation in response to low pH by enhancing nitrate uptake (Ye *et al*., 2021). Owing to this essential role in cellular organic acid and ion homeostasis, STOP1 is regulated at the protein level via ubiquitination, SUMOylation, and phosphorylation. The F-box protein REGULATION of AtAlMT1 EXPRESSION 1 (RAE1), together with its isoform RAE1 HOMOLOG 1 (RAH1), aids the ubiquitination of STOP1 and controls its degradation through the 26S proteasome (Fang *et al*., 2021). The small ubiquitin-like modifier (SUMO) E3 ligase SAP AND MIZ1 DOMAIN-CONTAINING LIGASE1 (SIZ1) and SUMO protease EARLY IN SHORT DAYS4 (ESD4) all coordinately modulate the abundance and activity of STOP1 by participating in its SUMOylation and deSUMOylation (Fang *et al*., 2020; Xu *et al*., 2021). Moreover, MAP KINASE 4 (MPK4) positively controls STOP1 accumulation through its phosphorylation, which weakens its interaction with RAE1 and downregulates its degradation (Zhou *et al*., 2023). Cytokinin is reportedly involved in STOP1-mediated proton toxicity resistance (Jiang *et al*., 2022). Nevertheless, the interconnection between the STOP1-mediated ion homeostasis pathway and abscisic acid (ABA) remains uncertain with respect to the underlying mechanism and interaction.

ABA is a critical phytohormone that plays key roles in normal plant growth and development, as well as in integrating stress signals and responses throughout the plant life cycle (Song *et al*., 2016). ABA is involved in multiple plant developmental stages, including embryonic maturation, seed dormancy and germination, seedling establishment, leaf senescence and abscission, and plant responses to various biotic and abiotic stress conditions (Chen *et al*., 2020; Cutler *et al*., 2010; Yoshida *et al*., 2019; Zhu, 2016). The major ABA signaling pathway consists of three core components: ABA receptors (PYRABACTIN RESISTANCE1 (PYR1)/PYR1-LIKE (PYL)/REGULATORY COMPONENTS OF ABA RECEPTOR (RCAR)), type 2C protein phosphatases (PP2Cs), and sucrose nonfermenting1 (SNF1)-related protein kinase 2s (SnRK2s) (Fujii *et al*., 2009; Ma *et al*., 2009; Park *et al*., 2009; Umezawa *et al*., 2010). When perceived by ABA receptors (PYR/PYL/RCAR), increased ABA levels can relieve the PP2Cs-mediated inhibition of downstream protein kinases, such as SnRK2s, leading to the phosphorylation of transcription factors (TFs) and altered expression of stress-responsive genes (Raghavendra *et al*., 2010; Wang *et al*., 2018). Particularly, the basic leucine zipper (bZIP) TF ABSCISIC ACID INSENSITIVE5 (ABI5), is an important regulator of ABA signaling. Specifically, ABI5 is activated by phosphorylated SnRK2s and enhances the expression of ABA-responsive genes (Yoshida *et al*., 2019). In turn, ABI5 can activate the expression of PP2Cs involving *ABI1* and *ABI2*, which function as a negative feedback in ABA signaling (Wang *et al*., 2019). Moreover, ABI5 can alter the expression of ABA receptor *PYLs* in response to exogenous ABA treatment during seed germination (Zhao *et al*., 2020). Further, the ABA receptor can be post-transcriptionally regulated. Meanwhile, the protein kinase CYTOSOLIC ABA RECEPTOR KINASE 1 (CARK1) can phosphorylate the ABA receptor and positively regulate the ABA response (Li *et al*., 2019; Zhang *et al*., 2018). ABA signaling can help plants to cope with low pH by regulating genes involved in aluminum (Al) tolerance and detoxification (Daspute *et al*., 2017; Hou *et al*., 2010; Ranjan *et al*., 2021). Thus, for example, in Arabidopsis, ABA induces the expression of *ALMT1*; however, an enhanced ABA-signaling mutant *abi1-1,* does not show changes in sensitivity to Al treatment (Kobayashi *et al*., 2013a). Conversely, *abi5* mutants show increased sensitivity to Al stress, but are independent of *ALMT1* and *multidrug and toxic compound extrusions* (*MATE*) expression (Fan *et al*., 2019). Similarly, the Al tolerance gene *Sensitive To Al Rhizotoxicity* (*OsSTAR1*) in rice is regulated by *STOP1 homolog* (*OsART1*) and members of the ASR (ABA-stress and ripening) gene family (Arenhart *et al*., 2014; Huang *et al*., 2009; Yamaji *et al*., 2009).In contrast, toxic metals can lead to the accumulation of ABA in plant tissues, such as root tips, and the stimulation of ABA-responsive gene expression (Ahmed *et al*., 2016; Hou *et al*., 2010; Sawaki *et al*., 2016; Vishwakarma *et al*., 2017). Nonetheless, to date, the mechanisms underlying ABA signaling and STOP1-mediated ion homeostasis remain largely unknown.

In this study, we observed that exogenous ABA treatment induced the expression of *RAE1*, leading to increased STOP1 degradation. Notably, *rae1* and *STOP1OE* plants exhibited enhanced sensitivity to exogenous ABA. This increased sensitivity was associated with the post-transcriptional upregulation of ABI5, a key TF involved in ABA signaling. Collectively, our findings suggest the existence of an ABA-responsive RAE1-STOP1 module that reduces sensitivity to exogenous ABA treatment in Arabidopsis by preventing the increase of ABI5.

## Results

### Exogenous ABA treatment induced *RAE1* expression

Publicly accessible transcriptome data in ePlants suggest that *RAE1* gene expression can be stimulated by ABA (Winter *et al*., 2007). We tested this hypothesis through real-time quantitative PCR and GUS (β-glucuronidase) activity assay. In particular, *RAE1* expression was significantly enhanced over a time course upon ABA treatment at 10 μM (**Figure 1A**). GUS activity was measured in the root of 2 d old seedlings of a stable *pRAE1:GUS* Arabidopsis transformation line. The GUS signal significantly increased after treatment with ABA for 3 h (**Figure 1B-C**). For long-term ABA treatment, from seed germination for up to 6 d, we observed enhanced GUS signals after ABA treatment at 1 μM (**Figure 1D**). These results demonstrate that ABA induced the expression of *RAE1* in Arabidopsis. Additionally, GUS-staining revealed that *RAE1* was mainly expressed in the roots and veins of *Arabidopsis thaliana*.

**Figure 1.**
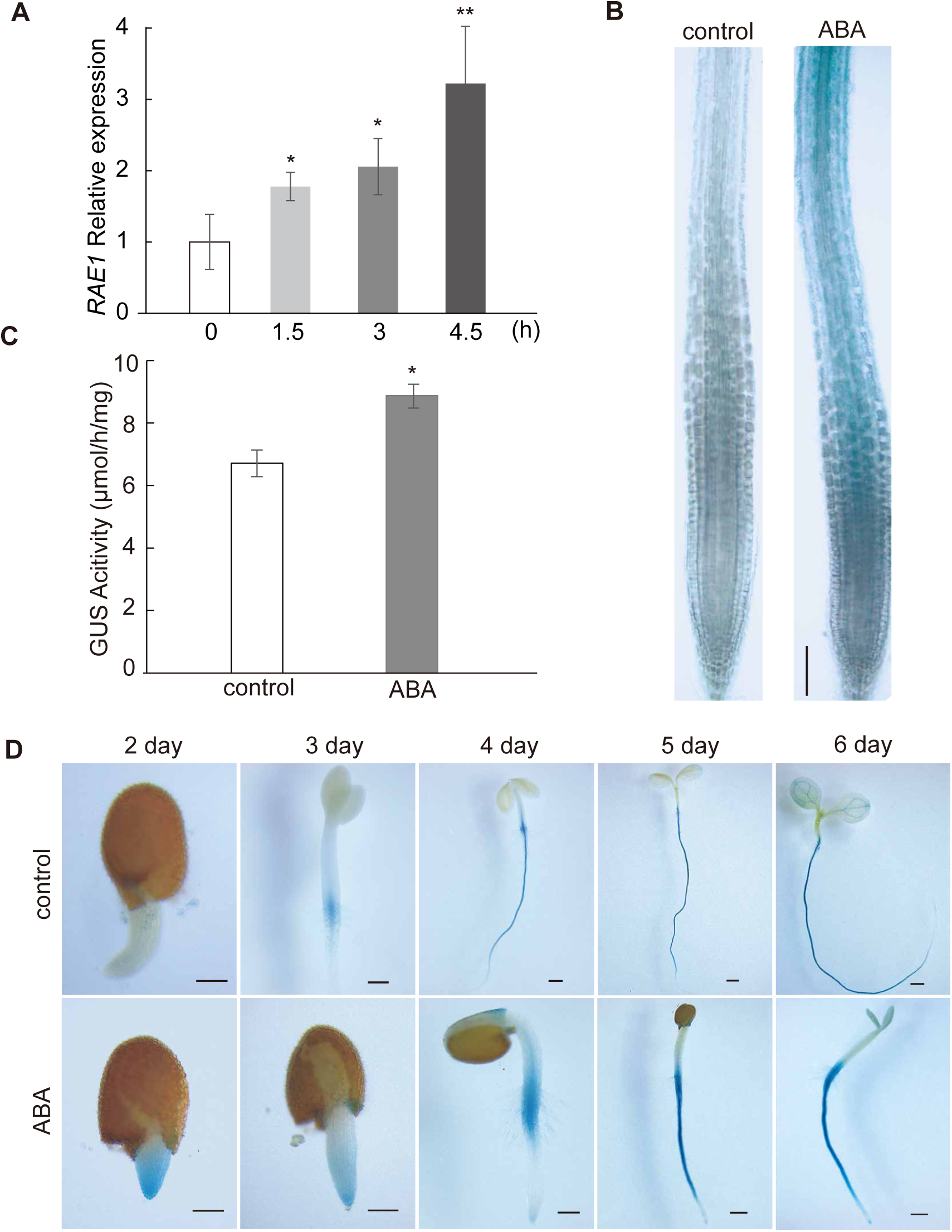
ABA induced *RAE1* gene expression in Arabidopsis. (**A**) The relative expression of *RAE1* in 7 d old plate grown Arabidopsis seedlings in response to ABA treatment are shown as column graphs. The level of *RAE1* transcripts after a time course of 10 μM ABA treatments (0, 1.5, 3, 4.5 h) were determined by qRT-PCR. (**B**) GUS-stained root tips from 5 d old seedlings of an Arabidopsis line with stabe expression of *pRAE1: GUS* were treated with or without 10 μM ABA treatment for 3 h for short-term ABA response analysis (scale bar=100 μm). (**C**) GUS activity was determined in 2 d old seedlings of the *pRAE1: GUS* line with or without 10 μM ABA treatment. (**D**) GUS stained 2 to 6 d old seed-lings of the *pRAE1: GUS* line with or without 1 μM ABA treatment from seeds (scale bar=200 μ m). Error bars represent the standard deviation for four biological replicates for qRT-PCR and GUS activity assay (* indicates significant difference (*P*<0.05), ** indicates highly significant difference (*P*<0.01), as per Student’s *t*-test).

### *rae1* mutant lines showed increased ABA sensitivity at the early seedling stage

To investigate whether *RAE1* is involved in the ABA response, we conducted an ABA sensitivity assay using three different *rae1* homozygous T-DNA insertion lines and their corresponding segregated wild type (WT) derived from the same F1 plants (**Figure S1A-C**). Although there was no change in seed germination in *rae1* mutant lines (**Figure S1D, E**), *rae1* seedlings showed a remarkably reduced proportion of greening cotyledons and fresh weight, compared with their corresponding WT counterparts **(Figure 2A-C**). To confirm that the ABA-sensitivity phenotype was due to a *RAE1* mutation, we generated two complementation lines (*RAE1 CM14-1* and *RAE1 CM2-2*) of the *rae1-b* mutant and performed the same ABA sensitivity assays. ABA sensitivity in the *rae1* mutant lines was restored to the level of the wild type after complementation (**Figure 2A-C, Figure S2**), indicating that *rae1* ABA-sensitive phenotype at the early seedling stage was indeed caused by a deficient *RAE1*. GUS-staining at the leaf development stage revealed that the expression of *RAE1* was mainly distributed in the veins (**Figure S3A, C**), but there was no apparent GUS-staining in the guard cells (**Figure S3B**). Therefore, no significant difference was observed in stomatal opening or water loss rate between the *rae1-b* mutant and WT under ABA treatment (**Figure S3D-F**). These results suggest that *RAE1* negatively regulated the ABA response, specifically at the early seedling stage.

**Figure 2.**
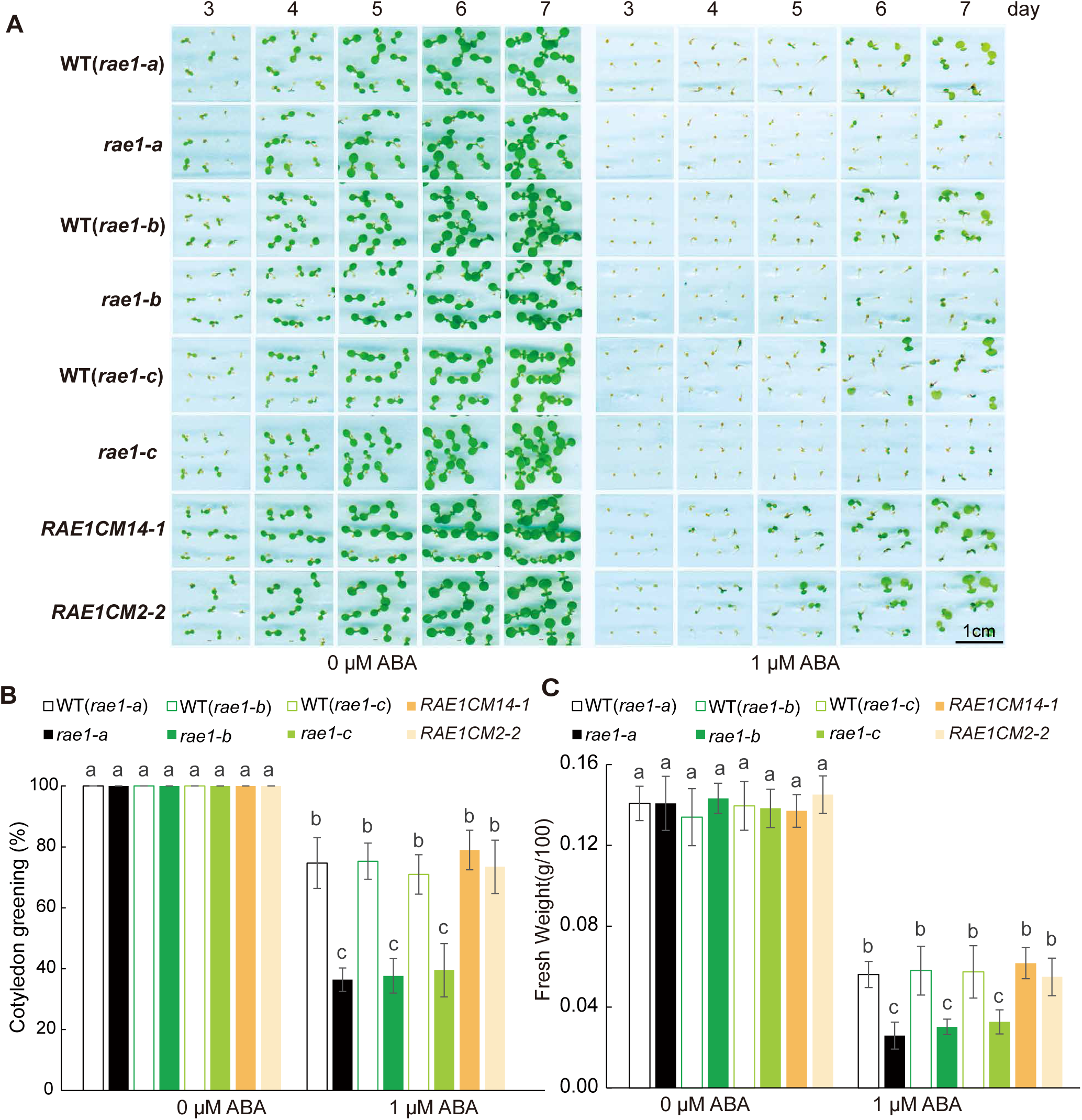
RAE1 loss of function leads to increased sensitivity to exogenous ABA treatment at the early seedling stage. (**A**) Seeds of *rae1-a, rae1-b, rae1-c*, their corresponding segregated wild type, and two complementation lines (*RAE1CM14-1* and *RAE1CM2-2*) were cultivated on half-strength MS medium with or without 1 μM ABA treatment. Photographs capturing representative images of seedlings aged between 3 and 7 d were taken daily and subsequently presented. (**B**-**C**) The proportion of 7 d old seedlings with cotyledon greening and fresh weight are shown as column graphs. Error bars represent the standard deviation for three repre-sentative plates. Grouping was determined by one-way ANOVA (*P*<0.05, Duncan’s multiple-range test).

### STOP1 mediated the ABA highly-sensitive phenotype in the early stage of *rae1* seedling establishment

We hypothesized that the increased sensitivity of *rae1* mutants to ABA is associated with the upregulation of STOP1, a known RAE1 substrate. To test this hypothesis, we performed an ABA sensitivity assay using two STOP1 overexpressing lines, *STOP1OE1* and *STOP1OE2*. As expected, both STOP1 overexpressing lines displayed significantly increased sensitivity to ABA (**Figure 3A-C**). A functional SCF (SKP1-CUL1-F-box) complex consists of SKP1 (ASK1), CULLIN 1 (CUL 1), RING-box 1 (RBX1), and F-box protein (Santner and Estelle, 2010), and the interaction between F-box protein and ASKs is essential for SCF-type E3 ligase activity. To ascertain the potential enhancement of the interaction between ABA-induced RAE1 and ASK, we co-transformed *RAE1-cEYFP* and *ASK1-nEYFP* into Arabidopsis protoplasts and subjected them to ABA treatment. The fluorescence signal, which represents the interaction between RAE1 and ASK1, was significantly enhanced after ABA treatment (**Figure S4A**, **B**). Furthermore, an *in vitro* ubiquitination assay of STOP1 was performed in protoplasts extracted from WT and *rae1* plants with or without ABA treatment. Briefly, *STOP1-2FLAG* and *UBQ10-3HA* were co-transformed into the protoplasts, STOP1 was enriched from the total protein using α-FLAG beads and the level of STOP1 ubiquitination was determined using a HA-antibody. This assay showed that ABA treatment increased STOP1 ubiquitination in the WT, but not in the *rae1-b* mutant (**Figure 3D**). Subsequently, we treated both the *rae1-b* mutant and WT seedlings with ABA and assessed the resulting alteration in STOP1 protein levels to determine whether exogenous ABA triggers STOP1 degradation in plants. STOP1 protein abundance was measured in 7 d old seedlings after 2 d of ABA treatment. We found a significant reduction in STOP1 protein abundance in the WT, whereas no significant change was observed in the *rae1-b* mutant (**Figure 3E, F**). Moreover, supplementation with the 26S proteasome inhibitor MG132 reduced the STOP1 decrease caused by ABA treatment in the WT plants. These results suggest that ABA-induced degradation of STOP1 by RAE1 mediates the ABA response in seedlings.

**Figure 3.**
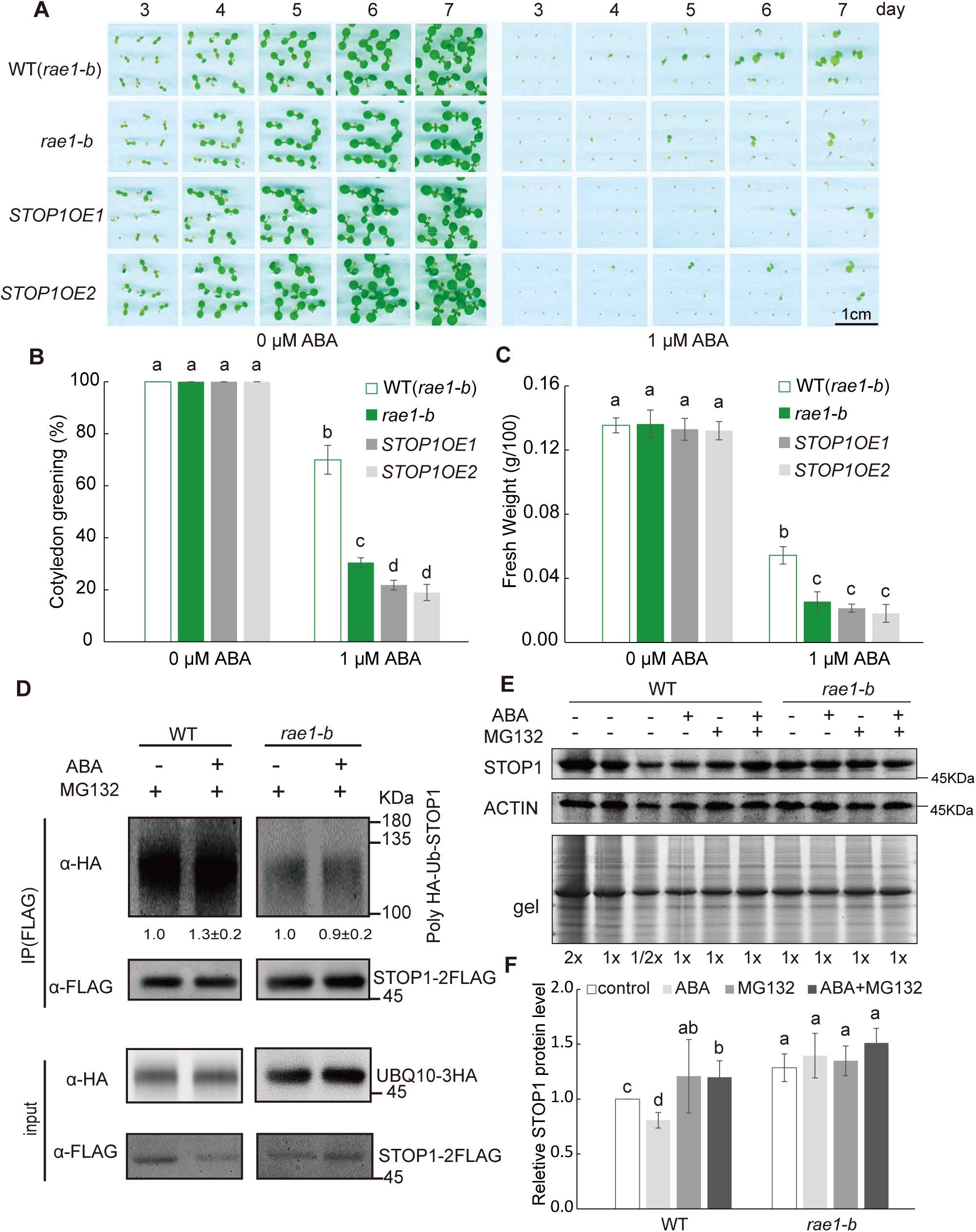
*STOP1* overexpression caused increased ABA sensitivity comparable to *rae1* at the early seedling stage. (**A**) Seeds of *rae1-b*, its corresponding segregated wild type, and two STOP1 overexpressing lines (*STOP1OE1* and *STOP1OE2*) were cultivated on half-strength MS medium with or without 1 μM ABA treatment. Photographs capturing representative images of seedlings aged between 3 and 7 d were taken daily and subsequently presented. (**B**-**C**) The proportion of 7 d old seedlings with cotyledon greening and fresh weight are shown as column graphs. Error bars represent the standard deviation for three representative plates. Grouping is determined by one-way ANOVA (*P*<0.05, Duncan’s multiple-range test). (**D**) ABA treatment increased the level of STOP1 ubiquitination in WT but not in *rae1-b* protoplasts. Two *pHBT* expression vectors containing either *35S:STOP1-2FLAG* or *UBQ10-3HA* were co-transformed into protoplasts extracted from WT or *rae1-b* Arabidopsis leaves and incubated with or without ABA/MG132 treatments. STOP1 was enriched from total protein using α-FLAG beads. Western blot analysis was performed to determine the abundance and ubiq-uitination of the enriched STOP1 using antibodies against HA/Flag. The numbers below the blotting indicate the relative level of STOP1 ubiquitination in either WT or *rae1-b* with or without ABA treatment (±SD for three biological replicates). (**E**) ABA treatment induced STOP1 reduction in WT but not in *rae1-b* mutants, which can be abolished by proteasome inhibitor MG132. 7 d old Arabidopsis seedlings grown on half-strength MS medium were transferred to new plates with or without 10 μM ABA and grown for another 2 d. Seedlings under control and ABA treatment were treated with or without 50 μM MG132 for another 6 h. Western blot of SDS-PAGE separated total protein was performed using an antibody against STOP1. Actin blotting and Coomassie blue stained gel were used for equal loading (**F**) STOP1 signal from blotting was determined using ImageJ and is represented in column graphs. Grouping was determined by Student’s *t*-test (*P* < 0.05). Error bars represent the standard deviation for three biological replicates.

### Increased sensitivity of the *rae1* mutant to ABA was mediated by ABI5 upregulation

STOP1 can regulate the cellular organic acid and ion status by activating the expression of *ALMT1*, *CIPK23* and *GDHs* which participate in malate secretion, ion transport, and metabolism associated to pH regulation (Sadhukhan *et al*., 2019; Sadhukhan *et al*., 2021), possibly affecting ABA absorption. To investigate whether the increased sensitivity of *rae1* mutants to ABA was due to increased absorption of exogenous ABA through transporters regulated by STOP1, we used LC-MS to measure the ABA content in *rae1-b* and the corresponding WT after ABA treatment. No significant difference in ABA content was observed between *rae1-b* and the WT after ABA treatment (**Figure S5A**). Other phytohormones, including indole-3-acetic acid (IAA), salicylic acid (SA), and jasmonic acid (JA), were also measured but no significant differences were observed (**Figure S5B, C, D**). To explore whether the increased ABA sensitivity observed in *rae1* mutants was influenced by *ALMT1*, which is a target gene of STOP1, we tested the ABA sensitivity of *ALMT1* overexpressing and mutant lines. The results showed that the sensitivity to ABA in both the *ALMT1* overexpressing lines and *almt1* mutant lines was similar to that in the WT (**Figure S6A, B**).

To characterize the enhanced ABA-sensitive *rae1* phenotype, we performed RNA-seq analysis on *rae1-b* and WT seedlings after ABA treatment. All the genes measured in the transcriptomic data are presented as volcanic plots (**Figure S7A**). A total of 22,629 genes were identified among these data, 211 which were marked in red for significant up-regulation in *rae1-b* mutants, while 206 genes were marked in blue for significant down-regulation in *rae1-b* mutants. Notably, the activation and repression of ABA-positive and ABA-negative regulatory factors were prevalent in *rae1-b* mutants (**Figure 4A**). Surprisingly, only four genes in the core ABA transduction signals (ABA receptors, PP2Cs, and SNRK2s) showed significant differences. Among them, the ABA receptors *PYL10* and *SNRK2.9* decreased, while two PP2Cs genes, *ABI1* and *ABI2*, increased slightly at the transcriptional level (**Figure 4B**). This finding indicated that ABA core signal-transduction changed little or was slightly inhibited in *rae1* under ABA treatment. Why are *rae1* mutants more sensitive to ABA? Although ABI5 transcript levels did not differ between *rae1-b* mutants and the WT, ABI5 downstream genes, such as *EARLY METHIONINE-LABELLED 6* (*EM6*), *RESPONSIVE TO ABA 18* (*RAB18*), *RESPONSIVE TO DESICCATION 29B* (*RD29B*), and *MOTHER OF FT AND TFL1* (*MFT*), were significantly upregulated in *rae1-b* compared with the WT (**Figure 4C**), indicating a potential post-transcriptional upregulation of ABI5 in *rae1*. To test this hypothesis, we used an antibody against ABI5 to determine and compare the abundance of ABI5 in *rae1-b*, *stop1-2* and *STOP1OE1*. Compared with the WT, the abundance of the ABI5 protein was higher in *rae1-b* and *STOP1OE1*, and lower in *stop1-2*, regardless of ABA treatment (**Figure 5A, B**). Furthermore, we generated an *abi5-8::rae1-b* double mutant by crossing *rae1-b* with *abi5-8* and performed an ABA sensitivity assay (**Figure 5C-E**). The introduction of *abi5-8* significantly reduced ABA sensitivity in *rae1-b*. In contrast, the introduction of *cark1*, a kinase that activates the ABA receptors PYL8 and PYR1 by phosphorylation (Zhang *et al*., 2018), did not diminish the ABA sensitivity phenotype in *rae1-b*. This observation implies that the ABA-sensitive trait in *rae1* was likely influenced by ABI5, prompting the question whether ABI5 is a target of RAE1-mediated ubiquitination. STOP1 can possibly interact with ABI5 to modulate its stability and functionality. However, yeast two-hybrid and Bimolecular fluorescence complementation (BiFC) experiments suggested no direct interaction between ABI5 and either RAE1 or STOP1 (**Figure S8A, B**).

**Figure 4.**
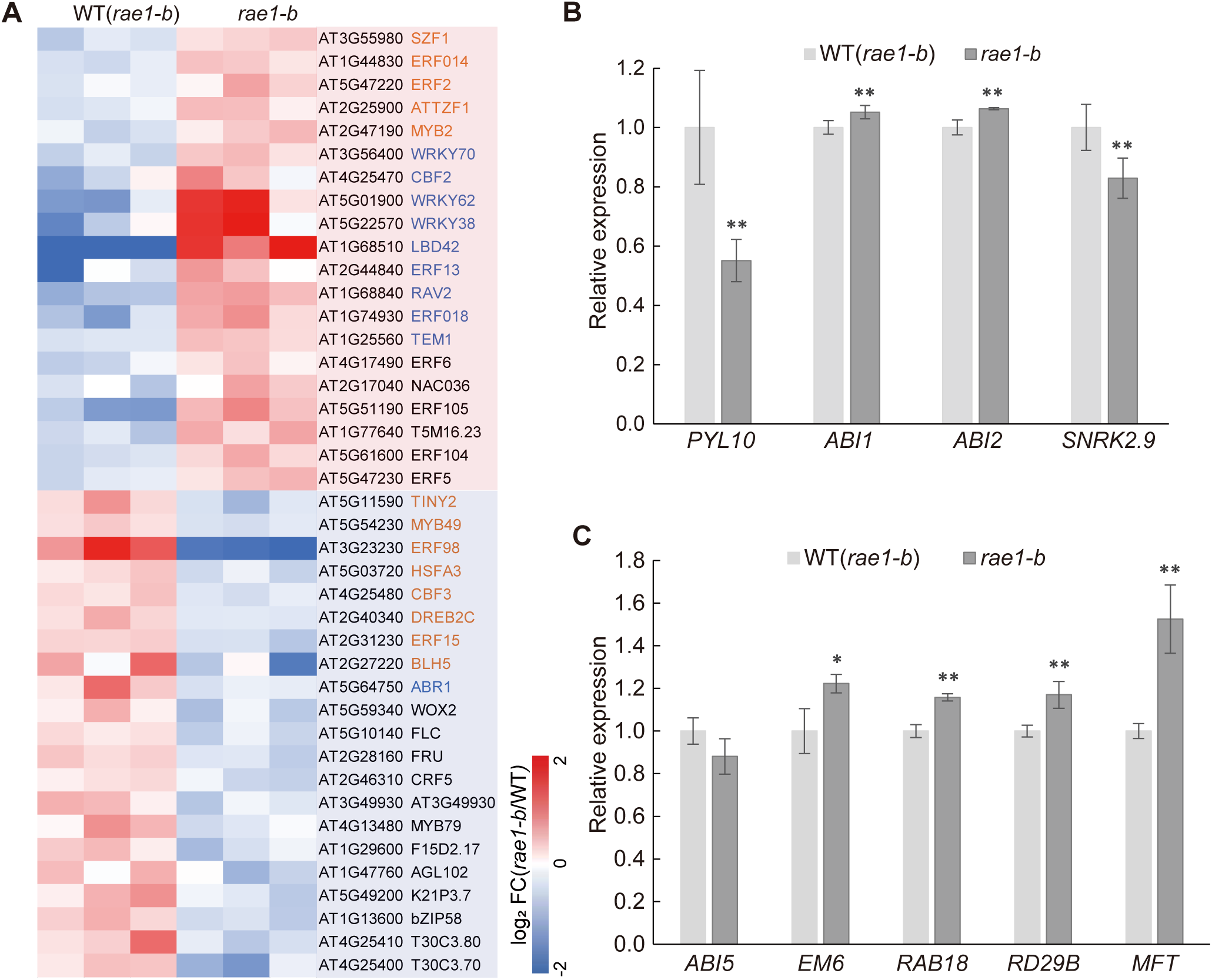
Upregulation of ABI5 target genes in *rae1-b*. (**A**) Heatmap showing transcription factors exhibiting significant changes (fold change >1.5 or <-1.5, *P*<0.05) in *rae1-b* compared to WT. Transcription factors characterized as ABA-positive regulators are high-lighted in red font, while negative regulators are highlighted in blue font. (**B**) Column graphs representing ABA receptors and core signaling factors, including *PYL10* (ABA receptor), *ABI1* (PP2C), *ABI2* (PP2C), and *SNRK2.9*, showing significant changes. (**C**) Transcript levels of *ABI5* target genes (*EM6*, *RAB18*, *RD29B*, *MFT*) and *ABI5* are shown as column graphs. Error bars indicate the standard deviation for three biological replicates (* indicates *P* < 0.05, ** indicates *P* < 0.01, as per Student’s *t*-test).

**Figure 5.**
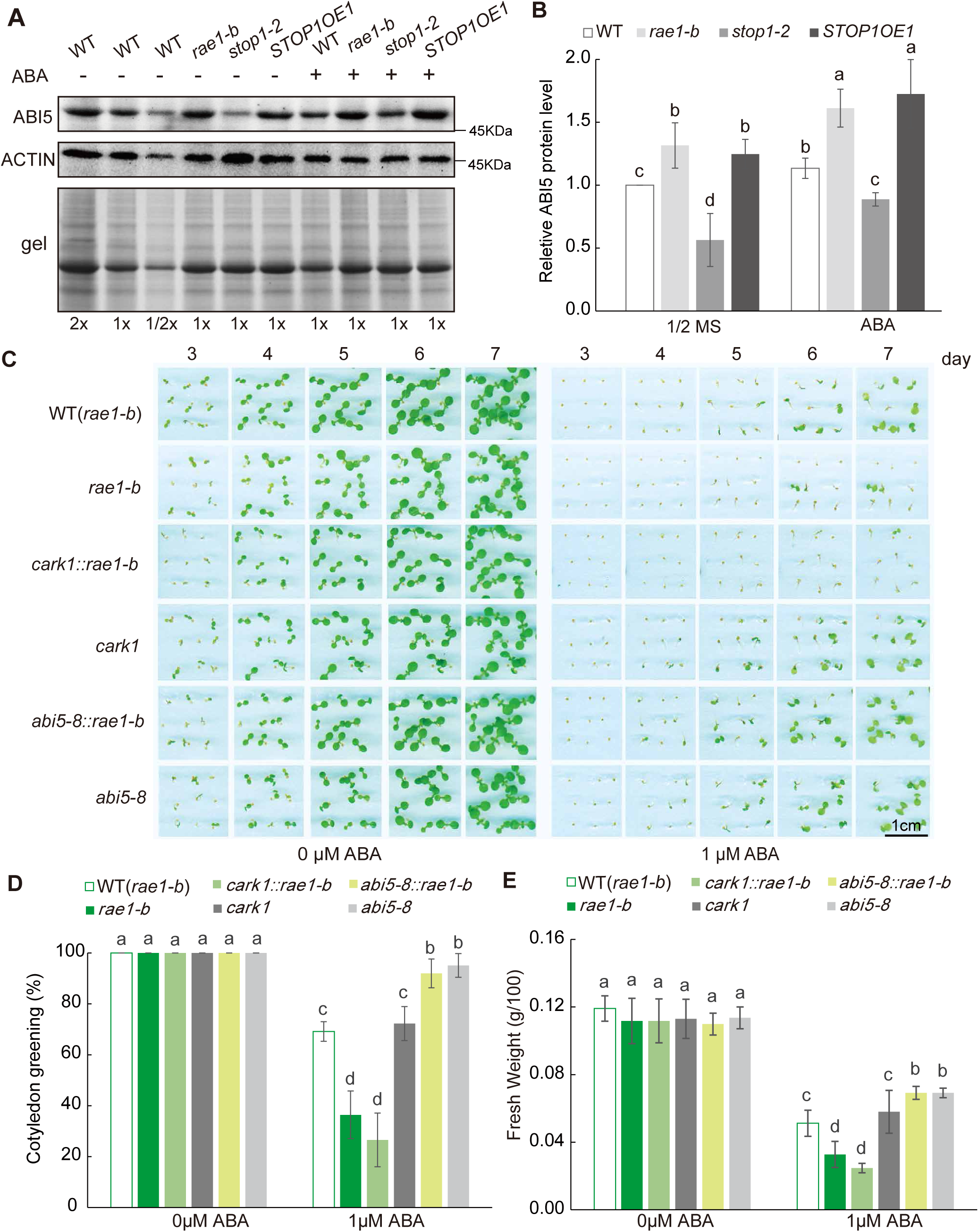
Heightened ABA sensitivity in *rae1* correlates with ABI5, not CARK1. (**A**) The abundance of ABI5 protein increased in *rae1-b* and *STOP1OE1*, but decreased in *stop1-2*, regard-less of ABA treatment. 5 d old Arabidopsis seedlings (WT, *rae1-b*, *stop1-2*, and *STOP1OE1*) were transferred to fresh plates with or without 10 μM ABA and grown for 20 h. Total protein extracted was separated by SDS-PAGE, followed by Western blot analysis using an ABI5 antibody. Actin blotting and Coomassie blue staining ensured equal loading. (**B**) ABI5 signals from blotting were normalized to WT and are presented as column graphs. Error bars represent the standard deviation for four biological replicates. (**C**) Mutant lines (*rae1-b*, *cark1*, *abi5-8*, *abi5-8::rae1-b*, and *cark1::rae1-b*) and WT were grown on half-strength MS medium with or without 1 μM ABA treatment. Daily photographs of seedlings aged between 3 and 7 d are shown (**D**-**E**) Column graphs depict the proportion of 7 d old seedlings with cotyledon greening and their fresh weight. Error bars represent the standard deviation for three representative plates. Grouping was determined by one-way ANOVA (*P*<0.05, Duncan’s multiple-range test).

## Discussion

The SCF-type E3 ligase F-box protein RAE1 downregulates STOP1 levels by enhancing its ubiquitination and subsequent degradation. Further, *RAE1* expression is induced by Al treatment and is dependent on STOP1 (Zhang *et al*., 2019). However, whether phytohormones regulate the RAE1-STOP1 module for cellular ion homeostasis or other responses remains unclear. In this study, we characterized RAE1 as an ABA-responsive mediator that contributes to lower ABA sensitivity associated with STOP1 at the early seedling stage. Indeed, *RAE1* expression was induced by exogenous ABA treatment and was mainly distributed in the vein tissues (**Figure 1C, D**). Consistently, exogenous ABA treatment enhanced the expression of *RAE1* and the formation of a functional SCF complex that ubiquitinated STOP1 and lowered its abundance at the early seedling stage (**Figure 3, Figure S4**). Moreover, *rae1* mutants exhibited increased ABA sensitivity during early seedling formation (**Figure 2**). Further, *STOP1* overexpressing lines were similar to the *rae1* lines in terms of ABA sensitivity (**Figure 3A-C**), suggesting that their increased ABA sensitivity was associated with the RAE1-STOP1 module. Furthermore, ABA and other phytohormones remained at the same level in *rae1* as in WT seedlings (**Figure S5**), indicating that the absorption of exogenous ABA or other phytohormones was not the cause. To determine whether this was associated with known STOP1 target genes, we further investigated *ALMT1* for its involvement in ABA sensitivity (**Figure S6**). However, the *almt1* knockout and overexpressing lines were similar to the WT plants under ABA treatment, indicating that the *rae1* ABA-sensitive phenotype cannot be explained by the known STOP1-ALMT1 pathway.

Additionally, we conducted a transcriptome analysis of *rae1-b* and compared it with that of the WT under ABA treatment to determine what genes participating in the ABA response are regulated by RAE1 (**Figure 4A, Figure S7A**).Changes in the transcript levels of ABA-positive and-negative regulatory factors were prevalent in *rae1-b* (**Figure 4A**), suggesting that the strong ABA response in *rae1-b* was a combined effect of positive and negative regulatory pathway factors in ABA signaling. This finding was consistent with previous reports suggesting that ABI5 not only functions as an important regulator of ABA signaling, but can also activate the expression of *ABI1* and *ABI2,* and change the expression pattern of ABA receptor PYLs, which function as a feedback of central ABA signaling (Collin *et al*., 2021; Wang *et al*., 2019; Zhao *et al*., 2020). Notably, the ABA receptor *PYL10* and the positive regulator *SNRK2.9* decreased at the transcript level, whereas two negative regulator PP2Cs genes, *ABI1* and *ABI2*, increased at the transcript level (**Figure 4B**). These findings suggest a potential upregulation of ABI5 in *rae1-b*. While there was no change in *ABI5* transcript levels in the *rae1-b*, a significant increase in the transcript levels of ABI5 target genes was observed (**Figure 4C**). As expected, ABI5 protein levels increased in both the *rae1* mutant and *STOP1* overexpressing lines, but decreased in *stop1-2* (**Figure 5A, B**). Moreover, the ABA sensitive phenotype *rae1-b* was fully restored by the ABI5 null allele *abi5-8* (**Figure 5C-D**). Altogether, these data indicate that the *rae1-b* ABA sensitive phenotype was by the ABI5-mediated ABA responses.

However, a direct interaction between ABI5 and RAE1 or STOP1 was not demonstrated. Moreover, the exact mechanism by which STOP1 regulates the increase in ABI5 protein remains unclear. However, Gene Ontology (GO) analysis revealed a substantial extent of upregulation of transporters and ubiquitin-related genes in the SCF complex of the *rae1-b* mutant (**Figure S7B, C**). Among them, several F-box-related genes were significantly upregulated (**Figure S7D**), particularly the homologous gene *RAH1*, which is closely associated with *RAE1*. This wide-scale transcriptome response to RAE1 defects suggests the presence of a complicated network that maintains cellular ion homeostasis through STOP1 and its crosstalk with other regulators of plant signaling involving ABA. A working model of RAE1 as an ABA-responsive factor that attenuates ABA signaling is schematized in (**Figure 6**). This working model proposes a link between exogenous ABA treatment and STOP1 through RAE1 and ABI5, suggesting that the STOP1 protein level must be precisely regulated to coordinate stress resistance and growth.

**Figure 6.**
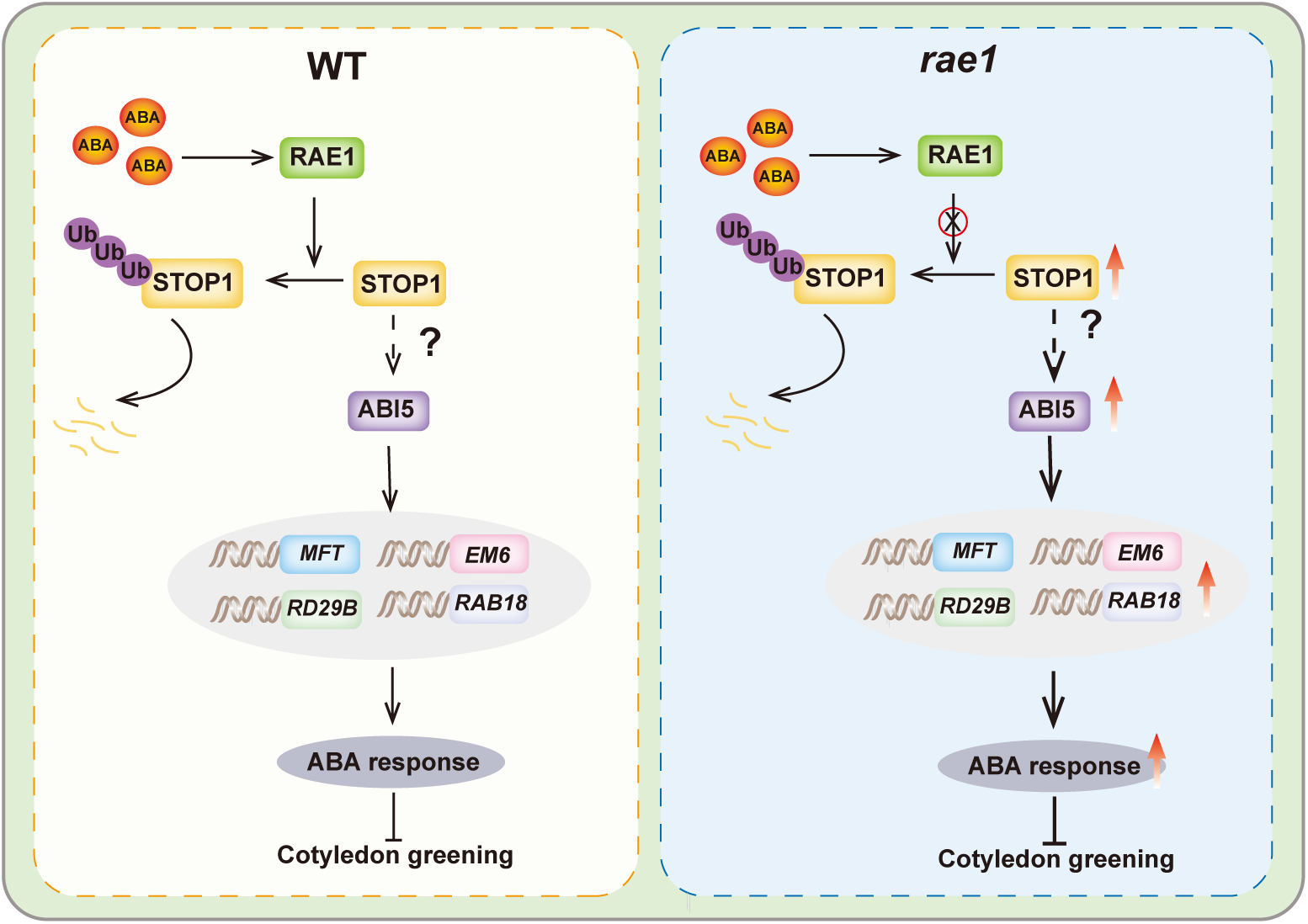
RAE1 functions as a responsive mediator to reduce ABA sensitivity through preventing ABI5 upregulation mediated by STOP1 at the early seedling stage. Exogenous ABA treatment can induce the expression of Arabidopsis *RAE1*, which contributes to STOP1 ubiquitination and degradation in the wild-type. However, in *rae1*, the lack of RAE1 associated degradation led to STOP1 accumulation and contributed to ABI5 upregulation via an unknown mediator. ABI5 upregulation stimulated the expression of target genes involved in *MFT*, *EM6*, *RAB18*, and *RD29B* and contributed to early seedling ABA sensitivity. Thus, RAE1-STOP1 serves as a response module to reduce ABA sensitivity by preventing the increase of ABI5 at the early seedling stage in Arabidopsis.

To further validate this proposed model, we tested the ABA sensitivity of the *stop1* mutant alongside *rae1-b* and compared them with wild type plants (**Figure S9A, B**). While *rae1-b* exhibits the highest sensitivity to exogenous ABA treatment among the three lines, we expected the *stop1* mutant to show lower ABA sensitivity due to the reduction of ABI5 abundance (**Figure 5A-B**). Surprisingly, the *stop1* mutant exhibited a slightly heightened sensitivity to exogenous ABA treatment compared to wild type plants, despite a clear reduction in ABI5 abundance. This suggests that exogenous ABA-induced RAE1-STOP1 interaction can reduce ABI5 abundance but not sufficiently to lower ABA sensitivity in wild type plants. Positive and/or feedback responses to exogenous ABA treatment in the *stop1* mutant line may explain this observation. Additionally, since STOP1 activates the transcription of RAE1 (Zhang *et al*., 2019), reduced RAE1 levels in the *stop1* mutant may lead to the accumulation of other potential, uncharacterized targets that contribute to ABA sensitivity, warranting further investigation in future studies.

## Conclusion

ABA induced *RAE1* expression, which in turn promoted STOP1 degradation. However, when *RAE1* was mutated or *STOP1* was overexpressed in Arabidopsis, plants showed increased sensitivity to exogenous ABA treatment, which was associated with the upregulation of ABI5, a key transcription factor involved in ABA signaling. Altogether, our findings suggest that the RAE1-STOP1 module was responsive to ABA and attenuated sensitivity to exogenous ABA sensitivity by regulating ABI5.

## Methods

### Plant Materials and Growth Conditions

In this study, we used *Arabidopsis thaliana* plants from the Columbia (*Col-0*) genetic background. T-DNA insertion lines *rae1-a* (SAIL_736_F09), *rae1-b* (SAIL_1053_H10), and *rae1-c* (SALK_103708) were obtained from the Nottingham Arabidopsis Stock Centre (NASC). Seeds of *abi5-8* (SALK_013163C) were obtained from Arashare. Double mutants *abi5-8::rae1-b* and *cark1::rae1-b*, were generated by crossing *rae1-b* with *abi5-8* and *cark1*. Other mutant lines, *pRAE1:GUS*, *stop1-2* (SALK_114108), *rae1-1* (G499A), *STOP1OE* (*OE1* and *OE2*), *cark1* (SALK_113377), *35s ALMT1-34*, *35s ALMT1-35* and *almt1-ko* (SALK_00962) were reported previously (Kobayashi *et al*., 2013b; Wang *et al*., 2021; Zhang *et al*., 2018; Zhang *et al*., 2019).A DNA fragment harboring a 2.4 kb *RAE1* promoter and a 4 kb genome sequence were cloned into the *pCAMBIA1301* vector. The resulting vector was used to transform into *rae1-b* and T3 homozygous plants for complementation. All seeds were surface-sterilized by soaking with bleach (5%, v/v) for 5 min and then washed five times with sterile water. Subsequently, sterilized seeds were placed in petri dishes on half strength Murashige and Skoog (MS) solid medium (1% sucrose and 0.8% agar) and incubated at 4 °C for 48 h. Seedlings were then grown in an incubator under long-day conditions (16 h light at 23 °C / 8 h dark at 18 °C). After 7 d of growth in the petri dishes, the seedlings were transferred to soil under the same long-day conditions. Light intensity for plant growth was 120-150 µmol m^-2^ s^-1^. Light was provided by T5 tubular LED lights (Leishi, China) at a color temperature of 6500K.

### GUS Histochemical analysis to determine *RAE1* expression

The GUS staining and experimental procedures were performed as previously described (Zhang *et al*., 2019). 5-d old seedlings or germinated seeds were harvested and treated with or without ABA for 3 h and incubated in freshly prepared GUS staining solution for an additional at 37 °C in darkness. Seedlings or seeds were washed three times with 70% ethanol until no green color was visible. Stained seedlings or germinated seeds were observed under a stereomicroscope (Leica, M165FC). Stained roots were observed under a fluorescence microscope (Leica, DFC420C). Root tissue was ground into a fine powder under liquid nitrogen, and then GUS activity was determined using a β-glucuronidase kit (geruisi, G0579F) according to manufacturer instructions.

### Bimolecular fluorescence complementation (BiFC) assay in Arabidopsis protoplasts

The coding sequences of *RAE1* and *ASK1/ABI5* were cloned into *pSAT-cEYFP* and *pSAT-nEYFP* vectors, respectively. The resultant *RAE1-cEYFP* and *ASK1/ABI5-nEYFP* were co-transformed into Arabidopsis protoplasts with or without 10 μM ABA treatment. Protoplast preparation and transformation were performed as described previously (Yoo *et al*., 2007). Protoplasts were prepared from the fourth, fifth, sixth, and seventh true leaves of 3-4 week old soil-grown Arabidopsis plants. Transfected protoplasts were incubated for 16 h before the YFP signal was assayed using a Zeiss.LSM710 confocal microscope (excitation wavelength at 488 nm and emission wavelength at 500–530 nm). Images were acquired using a 40× lens with a pinhole diameter of 1 airy unit (corresponding to an optical slice of 4.37 μm). Confocal images were further processed using the ImageJ software for quantification.

### STOP1 ubiquitination assay in Arabidopsis protoplasts

Protoplasts were derived from the fourth, fifth, sixth, and seventh true leaves of 3-4-week-old soil-grown wild-type (WT) and *rae1-b* Arabidopsis plants. For *in vitro* translation, two pHBT expression vectors containing either *35S:STOP1-2FLAG* or *UBQ10-3HA* were co-transformed into protoplasts and subsequently incubated for 20 h with or without ABA treatment at 20 μM. Following the incubation period, protoplasts were subjected to treatment with MG132 for an additional 6 h before total protein extraction using an extraction buffer containing 50 mM Tris (pH 7.5), 150 mM NaCl, 0.1% sodium dodecyl sulfate (SDS), 1% Triton X-100, 1 mM EGTA, 1 mM DTT, 1× complete protease inhibitor cocktail, and 1 mM PMSF. Protein STOP1 was selectively enriched from the total protein extract using Anti-FLAG® M2 Magnetic Beads (Sigma-Aldrich, M8823). Subsequently, the ubiquitination status of STOP1 was assessed using an anti-HA antibody (Proteintech, 66006-2-Ig). Protoplast preparation and transformation procedures were performed as previously described (Yoo *et al*., 2007).

### Western blot assay

Abundance of the STOP1 protein was determined in 9-d old WT and *rae1-b* mutant plants. Briefly, 7-d old seedlings grown on half-strength MS solid-medium plates were transferred to new half-strength MS plates with or without 10 μM ABA for another 2 d. Control and ABA treated seedling were transferred to half strength liquid MS medium with or without 50 µM MG132 and incubated for 6 h. Seedlings (0.1 g fresh weight) were homogenized to a fine powder under liquid nitrogen before protein extraction using four volumes of extraction buffer containing 50 mM Tris pH 7.5, 150 mM NaCl, 0.1% sodium dodecyl sulfate (SDS), 1% Triton X-100, 1 mM EGTA, 1 mM DTT, 1× complete protease inhibitor cocktail, and 1 mM PMSF. Total protein in the extraction buffer was centrifuged at 12,000[g for 10 min at 4 °C to remove cell debris. The supernatant was collected, and protein concentration was measured using the Amidoblack protocol for equal loading in SDS–PAGE. Proteins were blotted to a PVDF membrane (0.45 μm, Millipore) and incubated in a rabbit polyclonal antibody against STOP1 (ABclonal, A21044 with 1:500 dilution). Horseradish peroxidase (HRP) anti-rabbit IgG (H+L) (Proteintech, SA00001 at 1:2,000 dilution) was added for visualization.

ABI5 protein abundance was determined in 6 d old Arabidopsis seedlings (WT, *rae1-b*, *stop1-2* and *STOP1OE1*). 5 d old half-strength MS plate-grown seedlings were transferred to new half-strength MS plates with or without 10 μM ABA for another day. Total protein extraction and western blot were performed as previously described, using an antibody against ABI5 (Agrisera, AS121863, 1:1000 dilution).

### Cotyledon greening assays

Sterilized seeds were sown on half strength MS plates with or without 1 μM ABA and vernalized at 4 °C for 2 d. Plates were transferred to a growth cabinet for long-term cultivation under long day conditions (16 h of light at 23 °C, followed by 8 h of darkness at 18 °C). The percentage of seedlings with greening cotyledons (emergence of visibly green cotyledons) was determined after 7 to 9 d.

### Measurement of stomatal apertures

Stomatal aperture of WT and *rae1-b* plants was measured following a previously reported method (Cheong *et al*., 2007). Leaves were collected from 3-week-old soil-grown Arabidopsis plants. Leaves (adaxial surface-up, abaxial surface-down) were soaked for 3 h in 10 mM MES-KOH buffer (pH 6.15) containing 50 mM KCl and 10 µM CaCl_2_ under continuous light at 22 °C. Leaves were then transferred to the same solutions with or without 10 µM ABA and incubated for an additional 3h. The abaxial epidermis of the leaves was sliced and placed on glass slides. Advanced upright fluorescence microscopy (OLYMPUS,BX63) was used for visualization. Images of stomata from multiple leaves were captured and stomatal apertures were calculated using the ImageJ software. Three independent experiments were performed to ensure the accuracy and reliability of the results.

### Water loss measurement on detached leaves

Leaves (4^th^, 5^th^ and 6^th^ true leaves) were collected from 3-week old WT and *rae1-b* Arabidopsis plants. To prevent water loss, the leaves were stored in a weighing dish at room temperature to ensure that they were not exposed to direct sunlight or high humidity. The initial weight (W1) and real-time weight (W2) were recorded at 15 min intervals for 2 h. The water loss rate was calculated using the following formula: Water loss rate(W1-W2) / W1×100%.

### Phytohormone assay by LC-MS

9-d old plate grown WT and *rae1-b* seedlings were subjected to 10 μM ABA treatment in half strength MS liquid medium for 3 h. 50 mg seedlings frozen in liquid nitrogen frozen were ground to a powder using motor and pestle. Phytohormones were extracted using 500 μL extraction solution consisting of IPA (isopropanol), H_2_O, and HCl (2:1:0.002). Deuterated phytohormones were spiked in the extraction buffer as internal standard. After incubation for thirty minutes in extraction buffer, 1 ml of chloroform (CHCl_3_) was added and incubated for another thirty minutes. The same volume supernatant was dehydrated in nitrogen gas. Dehydrated samples were solubilized by 0.1 ml methanol and filtered by 0.1 μm column. Samples were injected to a UPLC-MS for phytohormone identification and quantification. UPLC I-Class Settings: Mobile phase is A: 0.05% acetic acid in H_2_O, B: 0.05% acetic acid in acetonitrile. The chromatographic column is poresell EC-120 3 μm 100mm, with a sample loading volume of 5 μL and a flow rate of 0.3 mL/min. The column temperature is 35 °C and the sample temperature is 15 °C. Q-Exactive MS Settings: Ion Source is HESI, Spray Voltage (-) is 3000, Capillary Temperature is 320, Sheath Gas is 30, Aux Gas is 10, Spare Gas is 5, Probe Heater Temp. is 350, S-Lens RF Level is 55, FULL MS-SIM: Resolution: 70,000, AGC target is 3e6, Maximum IT is 100, Scan range is 50 to 750 m/z.

### Transcriptomic analysis

9-d old WT and *rae1-b* Arabidopsis seedling were treated with half strength MS liquid medium containing 10 μM ABA for 3 h. 100 mg samples were frozen in liquid nitrogen. RNA was extracted from biological triplicates. RNA extraction, sample bank quality control and sequencing were completed by OE B.otech (Shanghai, China) as a service. The mRNA of poly (A) tail was enriched by Oligo (dT) magnetic beads. RNA integrity was assessed using the Agilent 2100 Bioanalyzer (Agilent Technologies, Santa Clara, CA, USA). Samples with RNA integrity Number (RIN)≥7 were analyzed. The libraries were constructed using TruSeq Stranded mRNA LT Sample Prep Kit (Illumina, San Diego, CA, USA) according to the manufacturer’s instructions. These libraries were then sequenced on Illumina sequencing platform (DNBSEQ-T7) and generated paired end readings of 125bp/150bp. Raw data (raw readings) were processed using Trimmomatic (Bolger *et al*., 2014). Remove readings containing poly-N and low-quality reads to obtain a clean reading. The clean reading is then mapped to the reference genome using hisat2 (Kim *et al*., 2015). The FPKM (Roberts *et al*., 2011) value of each gene was calculated using cufflinks (Trapnell *et al*., 2010), and the read counts of each gene were obtained by htseq-count (Anders *et al*., 2015). DEGs were identified using the DESeq (Anders and Huber, 2012) R package functions Estimate Size Factors and nbinomTest *P*-value < 0.05 with fold change >1.5 was set as the threshold for significantly differential expression. Hierarchical cluster analysis of DEGs was performed to explore the gene expression patterns.

### Yeast two-hybrid assay

The coding sequences of *ABI5* and *RAE1* with or without mutations were cloned into *pGADT7* (GAL-4 activation domain) and *pGBKT7* (GAL-4-binding domain) plasmids, respectively. The lithium acetate yeast transformation method was used to introduce the constructs into the yeast strain AH109 cells. Briefly, yeast cells were incubated for 6 h at room temperature in a 500 μL solution containing 600-800 ng vectors and 1 M LiAc, 1 M Tris-HCL 0.5 M EDTA (pH=8.0) and 45% PEG4000. The transformed yeasts were plated onto a nutrient-deficient medium for the selection of positive interactions.

## Open accessible data

RNA-seq data can be accessed through the following link https://dataview.ncbi.nlm.nih.gov/object/PRJNA938824?reviewer=teh30msot8veq6lfg25bqiodq0

## Acknowledgements

Professor Hiroyuki Koyama (Faculty of Applied Biological Sciences, Gifu University) is thanked for providing *almt1-ko* (SALK_009629) and *ALMT1-OE* seeds. Professor Yi Wang (College of Biological Sciences, China Agricultural University) for providing *STOP1OE1* and *STOP1OE2* seeds. Professor Yi Yang (College of Biological Sciences, Sichuan University) for providing *cark1* (SALK_113377) seeds. Professor Zhen Li (College of Biological Sciences, China Agricultural University) for assistance in the ABA contents assay in Arabidopsis.

## Author Contribution

LL designed the research; YQZ performed plant culture and biochemical experiments; MH and YYL participated in the WB and BiFC experiments. YQH and MMY participated in the Y2H experiments. NNW and CFH contributed plant mutant lines, vectors and agents. LL and YQZ contributing to the writing and revision of the article.

## Funding

This work was supported through funding by the National Natural Science Foundation of China (31970294), Tianjin Natural Science Foundation (19JCYBJC24100), and Open Research Fund of State Key Laboratory of Hybrid Rice (Wuhan University KF202201) to LL and the National Natural Science Foundation of China (32070317) to NNW.

## Competing Interests

The Authors declare that there are no competing interests associated with the manuscript.

## Supplemental Figures

**Figure S1. Characterization of three *RAE1* T-DNA insertion lines.**

**(A)** The position of three T-DNA insertion loci is annotated in *RAE1* diagrams. Exons are annotated as boxes and introns are annotated as solid lines. (**B**, **C**) Expression of *RAE1* was determined by qRT-PCR in mutant lines and normalized to WT. qRT-PCR primers were designed to measure the expression of N-terminal and C-terminal region. Primers are annotated in the *RAE1* diagram. Error bars represent the standard deviation for three biological replicates. (* indicates significant difference (*P*<0.05), ** indicates highly significant difference (*P*<0.01), as per Student’s *t*-test). (**D**, **E**) Arabidopsis mutant lines (*rae1-a*, *rae1-b*, *rae1-c*, *almt1-ko*) along with WT, overexpressing lines *35S:ALMT1-34* and *35S:ALMT1-35* were seeded on half-strength MS medium with or without ABA. Seed coat rupture and emergence of radicles served as criteria for germination counting. Error bars represent the standard deviation for three biological replicates. Each plate with 36 plants was treated as one biological replicate.

**Figure S2. Validation of two *RAE1* complement lines using RT-PCR.**

(**A**, **B**) Expression of *RAE1* (*UBQ1* as internal control) in WT, *rae1-b*, *RAE1cm1-14*, and *RAE1cm2-2* were determined by RT-PCR. Total RNA was extracted from 7 d old plate grown Arabidopsis seedlings. Error bars represent the standard deviation for three biological replicates. (* indicates significant difference (*P*<0.05), ** indicates highly significant difference (*P*<0.01), as per Student’s *t*-test).

**Figure S3. *rae1* mutant showed the same ABA responses as WT at leaf production stage.**

**(A)** A stable transformation line expressing *pRAE1:GUS* was used for *RAE1* tissue specific expression analysis. GUS staining was performed using 3 week old Arabidopsis plants. (**B**) The adaxial leaf surface was focused to visualize GUS signal in stomata and vein tissues. (**C**) True leaves (1-10) were detached and arranged according to their emergence. (**D**) Relative water loss curve over 2 h are shown. Error bars represent the standard deviation for sixteen Arabidopsis plants (Student’s *t*-test). (**E, F**) Stomatal aperture with or without 10 μM ABA treatment was measured under an advanced upright fluorescence microscopy. Error bars represent the standard deviation for 21-24 Arabidopsis stomata from three to five leaves. Grouping was determined by one-way ANOVA (*P*<0.05, Duncan’s multiple-range test).

**Figure S4. ABA enhanced the interaction between ASK1 and STOP1.**

**(A)** *RAE1-cEYFP* and *ASK1-nEYFP* vectors were transformed into Arabidopsis protoplasts by PEG treatment with or without 10 μM ABA treatment. BiFC signal, which represents the RAE1-ASK1 interaction, was observed in nuclear compartments under a confocal microscope. (**B**) The fluorescence intensity of the interaction signal in 11 protoplasts was determined and compared between the control and ABA treatment (scale bar=5 μm). (* indicates significant difference (*P*<0.05), ** indicates highly significant difference (*P*<0.01), as per Student’s *t*-test).

**Figure S5. Contents of phytohormones in *rae1-b* mutant and wild type.**

(**A-D**) 9 d old *rae1-b* and WT seedlings were treated with 10 μM ABA for 3 h. ABA, SA, IAA, and JA contents were measured by LC-MS as described in Methods. Error bars represent the standard deviation for five biological replicates (*P*<0.05, Student’s *t*-test).

**Figure S6. Cotyledon greening assay in ALMT1 associated lines under ABA treatment.**

**(A)** Arabidopsis mutant lines (*almt1-ko*, *rae1* mutants), WT and overexpressing line *35S ALMT1-OE* were grown on half-strength MS medium with or without 1 μM ABA treatment. Photographs capturing representative images of seedlings aged between 3 and 7 d were taken daily and subsequently presented. (**B-C**) The proportion of 7 d old seedlings with cotyledon greening and fresh weight are shown as column graphs. Error bars represent the standard deviation for three representative plates. Grouping was determined by one-way ANOVA (P<0.05, Duncan’s multiple-range test).

**Figure S7. Transcriptome analysis in *rae1* by RNA deep sequencing.**

**(A)** Log transferred fold change in transcripts of 22,629 genes between *rae1-b* and WT are shown as volcano plotting. 211 genes with significant increase in transcript level are highlighted in red. 206 genes with significant reduction in transcript level are highlighted in blue. (**B**) GO enrichment analysis was performed using differentially expressed coding genes (FC>2, *P*<0.05) as target, and all encoding genes as background. The top 30 enriched GO terms for biological process, cellular components and molecular function are shown. (**C-D**) Transporters and F-box proteins with significantly changes (FC>1.5 or FC<-1.5, *P*<0.05) in *rae1-b* are shown as heatmap. (**E**) Sequencing frequency intensity covering *RAE1* is shown in the WT and *rae1-b*.

**Figure S8. The yeast two hybrid and BiFC assays did not indicate an interaction between RAE1-ABI5 or STOP1-ABI5.**

**(A)** Yeast two hybrid vectors *AD-ABI5*, *BD-RAE1*, *BD-RAE1* (G167R) point mutation, and *BD-RAE1* (75aa-655aa) missing F-box or *AD-STOP1*, and *BD-ABI5*, were transformed into yeast to investigate ABI5 and RAE1 or STOP1 and ABI5 interaction. *AD-ASK1* and *BD-RAE1* were used as a positive control. Nutrient deficient medium lacking Lue, Trp or lacking Lue, Trp, His, and Ade were used for screening. (**B**) The coding sequence of *RAE1/STOP1* and *ASK1/ABI5/STOP1* was cloned into the BiFC vectors *pSAT-cEYFP* and *pSAT-nEYFP* respectively. The resultant *RAE1/STOP1-cEYFP* and *ASK1/ABI5/STOP1-nEYFP* were co-transformed into Arabidopsis protoplasts. BiFC signal which represents the RAE1-ASK1 interaction were observed in nuclear compartments under a confocal microscope (scale bar=5 μm). However, no interaction signal was present in RAE1 and ABI5 or STOP1 and ABI5 co-transformations.

**Figure S9. The *stop1* mutant line exhibits slightly heightened sensitivity to exogenous ABA treatment, but not comparable to *rae1-b*.**

**(A)** Seeds of *rae1-b*, *stop1-2*, and wild type were grown under the same conditions for nine days. **(B)** The green cotyledon rate was measured and presented as column graphs. Error bars represent standard deviations for three representative plates. Grouping was determined by one-way ANOVA (P<0.05, Duncan’s multiple-range test).

## Supplemental Data

**DataS1** RNA deep sequencing to determine transcriptional changes in *rae1-b*.

**DataS2** Transcription factors with significant changes in expression.

**DataS3** Transporters with significant changes in expression.

**DataS4** F-box genes with significant changes in expression.

## Notes

### Competing Interest Statement

The authors have declared no competing interest.

## Reference

Ahmed IM, Nadira UA, Cao FB, He XY, Zhang GP, Wu FB. 2016. Physiological and molecular analysis on root growth associated with the tolerance to aluminum and drought individual and combined in Tibetan wild and cultivated barley. Planta 243, 973–985.

Anders S, Huber WJH, Germany: European Molecular Biology Laboratory. 2012. Differential expression of RNA-Seq data at the gene level–the DESeq package. European Molecular Biology Laboratory 10, f1000research.

Anders S, Pyl PT, Huber W. 2015. HTSeq--a Python framework to work with high-throughput sequencing data. Bioinformatics 31, 166–169.

Arenhart RA, Bai Y, de Oliveira LFV, Neto LB, Schunemann M, Maraschin FD, Mariath J, Silverio A, Sachetto-Martins G, Margis R, Wang ZY, Margis-Pinheiro M. 2014. New Insights into Aluminum Tolerance in Rice: The ASR5 Protein Binds the Promoter and Other Aluminum-Responsive Genes. Molecular Plant 7, 709–721.

Balzergue C, Dartevelle T, Godon C, Laugier E, Meisrimler C, Teulon JM, Creff A, Bissler M, Brouchoud C, Hagege A, Muller J, Chiarenza S, Javot H, Becuwe-Linka N, David P, Peret B, Delannoy E, Thibaud MC, Armengaud J, Abel S, Pellequer JL, Nussaume L, Desnos T. 2017. Low phosphate activates STOP1-ALMT1 to rapidly inhibit root cell elongation. Nat Commun 8, 15300.

Bolger AM, Lohse M, Usadel B. 2014. Trimmomatic: a flexible trimmer for Illumina sequence data. Bioinformatics 30, 2114–2120.

Chen K, Li GJ, Bressan RA, Song CP, Zhu JK, Zhao Y. 2020. Abscisic acid dynamics, signaling, and functions in plants. Journal of Integrative Plant Biology 62, 25–54.

Cheong YH, Pandey GK, Grant JJ, Batistic O, Li L, Kim BG, Lee SC, Kudla J, Luan S. 2007. Two calcineurin B-like calcium sensors, interacting with protein kinase CIPK23, regulate leaf transpiration and root potassium uptake in Arabidopsis. Plant Journal 52, 223–239.

Collin A, Daszkowska-Golec A, Szarejko I. 2021. Updates on the Role of ABSCISIC ACID INSENSITIVE 5 (ABI5) and ABSCISIC ACID-RESPONSIVE ELEMENT BINDING FACTORs (ABFs) in ABA Signaling in Different Developmental Stages in Plants. Cells 10.

Cutler SR, Rodriguez PL, Finkelstein RR, Abrams SR. 2010. Abscisic Acid: Emergence of a Core Signaling Network. Annual Review of Plant Biology, Vol 61 61, 651–679.

Daspute AA, Sadhukhan A, Tokizawa M, Kobayashi Y, Panda SK, Koyama H. 2017. Transcriptional Regulation of Aluminum-Tolerance Genes in Higher Plants: Clarifying the Underlying Molecular Mechanisms. Front Plant Sci 8, 1358.

Fan W, Xu JM, Wu P, Yang ZX, Lou HQ, Chen WW, Jin JF, Zheng SJ, Yang JL. 2019. Alleviation by abscisic acid of Al toxicity in rice bean is not associated with citrate efflux but depends on ABI5-mediated signal transduction pathways. J Integr Plant Biol 61, 140–154.

Fang Q, Zhang J, Zhang Y, Fan N, van den Burg HA, Huang CF. 2020. Regulation of Aluminum Resistance in Arabidopsis Involves the SUMOylation of the Zinc Finger Transcription Factor STOP1. Plant Cell 32, 3921–3938.

Fang Q, Zhou F, Zhang Y, Singh S, Huang CF. 2021. Degradation of STOP1 mediated by the F-box proteins RAH1 and RAE1 balances aluminum resistance and plant growth in Arabidopsis thaliana. Plant J 106, 493–506.

Fujii H, Chinnusamy V, Rodrigues A, Rubio S, Antoni R, Park SY, Cutler SR, Sheen J, Rodriguez PL, Zhu JK. 2009. reconstitution of an abscisic acid signalling pathway. Nature 462, 660–U138.

Hou NN, You JF, Pang JD, Xu MY, Chen G, Yang ZM. 2010. The accumulation and transport of abscisic acid insoybean (Glycine max L.) under aluminum stress. Plant and Soil 330, 127–137.

Huang CF, Yamaji N, Mitani N, Yano M, Nagamura Y, Ma JF. 2009. A bacterial-type ABC transporter is involved in aluminum tolerance in rice. Plant Cell 21, 655–667.

Jiang F, Lyi SM, Sun T, Li L, Wang T, Liu J. 2022. Involvement of cytokinins in STOP1-mediated resistance to proton toxicity. Stress Biol 2, 17.

Kim D, Langmead B, Salzberg SL. 2015. HISAT: a fast spliced aligner with low memory requirements. Nat Methods 12, 357–360.

Kobayashi Y, Kobayashi Y, Sugimoto M, Lakshmanan V, Iuchi S, Kobayashi M, Bais HP, Koyama H. 2013a. Characterization of the Complex Regulation of AtALMT1 Expression in Response to Phytohormones and Other Inducers. Plant Physiology 162, 732–740.

Kobayashi Y, Lakshmanan V, Kobayashi Y, Asai M, Iuchi S, Kobayashi M, Bais HP, Koyama H. 2013b. Overexpression of AtALMT1 in the Arabidopsis thaliana ecotype Columbia results in enhanced Al-activated malate excretion and beneficial bacterium recruitment. Plant Signal Behav 8.

Li XY, Kong XG, Huang Q, Zhang Q, Ge H, Zhang L, Li GM, Peng L, Liu ZB, Wang JM, Li XF, Yang Y. 2019. CARK1 phosphorylates subfamily III members of ABA receptors. Journal of Experimental Botany 70, 519–528.

Ma Y, Szostkiewicz I, Korte A, Moes D, Yang Y, Christmann A, Grill E. 2009. Regulators of PP2C Phosphatase Activity Function as Abscisic Acid Sensors. Science 324, 1064–1068.

Park SY, Fung P, Nishimura N, Jensen DR, Fujii H, Zhao Y, Lumba S, Santiago J, Rodrigues A, Chow TFF, Alfred SE, Bonetta D, Finkelstein R, Provart NJ, Desveaux D, Rodriguez PL, McCourt P, Zhu JK, Schroeder JI, Volkman BF, Cutler SR. 2009. Abscisic Acid Inhibits Type 2C Protein Phosphatases via the PYR/PYL Family of START Proteins. Science 324, 1068–1071.

Raghavendra AS, Gonugunta VK, Christmann A, Grill E. 2010. ABA perception and signalling. Trends in Plant Science 15, 395–401.

Ranjan A, Sinha R, Lal SK, Bishi SK, Singh AK. 2021. Phytohormone signalling and cross-talk to alleviate aluminium toxicity in plants. Plant Cell Rep 40, 1331–1343.

Roberts A, Trapnell C, Donaghey J, Rinn JL, Pachter L. 2011. Improving RNA-Seq expression estimates by correcting for fragment bias. Genome Biol 12, R22.

Sadhukhan A, Enomoto T, Kobayashi Y, Watanabe T, Iuchi S, Kobayashi M, Sahoo L, Yamamoto YY, Koyama H. 2019. Sensitive to Proton Rhizotoxicity1 Regulates Salt and Drought Tolerance of Arabidopsis thaliana through Transcriptional Regulation of CIPK23. Plant and Cell Physiology 60, 2113–2126.

Sadhukhan A, Kobayashi Y, Iuchi S, Koyama H. 2021. Synergistic and antagonistic pleiotropy of STOP1 in stress tolerance. Trends in Plant Science 26, 1014–1022.

Santner A, Estelle M. 2010. The ubiquitin-proteasome system regulates plant hormone signaling. Plant Journal 61, 1029–1040.

Sawaki K, Sawaki Y, Zhao CR, Kobayashi Y, Koyama H. 2016. Specific transcriptomic response in the shoots of Arabidopsis thaliana after exposure to Al rhizotoxicity: - potential gene expression biomarkers for evaluating Al toxicity in soils. Plant and Soil 409, 131–142.

Sawaki Y, Iuchi S, Kobayashi Y, Kobayashi Y, Ikka T, Sakurai N, Fujita M, Shinozaki K, Shibata D, Kobayashi M, Koyama H. 2009. STOP1 Regulates Multiple Genes That Protect Arabidopsis from Proton and Aluminum Toxicities. Plant Physiology 150, 281–294.

Song L, Huang SSC, Wise A, Castanon R, Nery JR, Chen HM, Watanabe M, Thomas J, Bar-Joseph Z, Ecker JR. 2016. A transcription factor hierarchy defines an environmental stress response network. Science 354.

Trapnell C, Williams BA, Pertea G, Mortazavi A, Kwan G, van Baren MJ, Salzberg SL, Wold BJ, Pachter L. 2010. Transcript assembly and quantification by RNA-Seq reveals unannotated transcripts and isoform switching during cell differentiation. Nat Biotechnol 28, 511–515.

Umezawa T, Nakashima K, Miyakawa T, Kuromori T, Tanokura M, Shinozaki K, Yamaguchi-Shinozaki K. 2010. Molecular Basis of the Core Regulatory Network in ABA Responses: Sensing, Signaling and Transport. Plant and Cell Physiology 51, 1821–1839.

Vishwakarma K, Upadhyay N, Kumar N, Yadav G, Singh J, Mishra RK, Kumar V, Verma R, Upadhyay RG, Pandey M, Sharma S. 2017. Abscisic Acid Signaling and Abiotic Stress Tolerance in Plants: A Review on Current Knowledge and Future Prospects. Front Plant Sci 8, 161.

Wang K, He JN, Zhao Y, Wu T, Zhou XF, Ding YL, Kong LY, Wang XJ, Wang Y, Li JG, Song CP, Wang BS, Yang SH, Zhu JK, Gong ZZ. 2018. EAR1 Negatively Regulates ABA Signaling by Enhancing 2C Protein Phosphatase Activity. Plant Cell 30, 815–834.

Wang XJ, Guo C, Peng J, Li C, Wan FF, Zhang SM, Zhou YY, Yan Y, Qi LJ, Sun KW, Yang SH, Gong ZZ, Li JG. 2019. ABRE-BINDING FACTORS play a role in the feedback regulation of ABA signaling by mediating rapid ABA induction of ABA co-receptor genes. New Phytologist 221, 341–355.

Wang ZF, Mi TW, Gao YQ, Feng HQ, Wu WH, Wang Y. 2021. STOP1 Regulates LKS1 Transcription and Coordinates K(+)/NH(4)(+) Balance in Arabidopsis Response to Low-K(+) Stress. Int J Mol Sci 23.

Winter D, Vinegar B, Nahal H, Ammar R, Wilson GV, Provart NJ. 2007. An "Electronic Fluorescent Pictograph" browser for exploring and analyzing large-scale biological data sets. PLoS One 2, e718.

Xu J, Zhu J, Liu J, Wang J, Ding Z, Tian H. 2021. SIZ1 negatively regulates aluminum resistance by mediating the STOP1-ALMT1 pathway in Arabidopsis. J Integr Plant Biol 63, 1147–1160.

Yamaji N, Huang CF, Nagao S, Yano M, Sato Y, Nagamura Y, Ma JF. 2009. A zinc finger transcription factor ART1 regulates multiple genes implicated in aluminum tolerance in rice. Plant Cell 21, 3339–3349.

Ye JY, Tian WH, Zhou M, Zhu QY, Du WX, Zhu YX, Liu XX, Lin XY, Zheng SJ, Jin CW. 2021. STOP1 activates NRT1.1-mediated nitrate uptake to create a favorable rhizospheric pH for plant adaptation to acidity. Plant Cell 33, 3658–3674.

Yoo SD, Cho YH, Sheen J. 2007. Arabidopsis mesophyll protoplasts: a versatile cell system for transient gene expression analysis. Nature Protocols 2, 1565–1572.

Yoshida T, Christmann A, Yamaguchi-Shinozaki K, Grill E, Fernie AR. 2019. Revisiting the Basal Role of ABA - Roles Outside of Stress. Trends in Plant Science 24, 625–635.

Zhang L, Li X, Li D, Sun Y, Li Y, Luo Q, Liu Z, Wang J, Li X, Zhang H, Lou Z, Yang Y. 2018. CARK1 mediates ABA signaling by phosphorylation of ABA receptors. Cell Discov 4, 30.

Zhang Y, Zhang J, Guo J, Zhou F, Singh S, Xu X, Xie Q, Yang Z, Huang CF. 2019. F-box protein RAE1 regulates the stability of the aluminum-resistance transcription factor STOP1 in Arabidopsis. Proc Natl Acad Sci U S A 116, 319–327.

Zhao HY, Nie KL, Zhou HP, Yan XJ, Zhan QD, Zheng Y, Song CP. 2020. ABI5 modulates seed germination via feedback regulation of the expression of the ABA receptor genes. New Phytologist 228, 596–608.

Zhou F, Singh S, Zhang J, Fang Q, Li C, Wang J, Zhao C, Wang P, Huang CF. 2023. The MEKK1-MKK1/2-MPK4 cascade phosphorylates and stabilizes STOP1 to confer aluminum resistance in Arabidopsis. Mol Plant 16, 337–353.

Zhu JK. 2016. Abiotic Stress Signaling and Responses in Plants. Cell 167, 313–324.

